# Dissociable Contributions of the Right Dorsolateral Prefrontal and Posterior Parietal Cortices to Automatic Alcohol Approach Tendencies

**DOI:** 10.64898/2026.04.19.719365

**Authors:** Adarsh K. Verma, Adith Deva Kumar, Usha Chivukula, Neeraj Kumar

## Abstract

Compulsive alcohol intake is often sustained by approach tendencies that automatically orient individuals toward alcohol-related cues, even when they intend to avoid them. Converging evidence implicates the frontoparietal regions in regulating such automatic behavior; however, the specific contributions of the right dorsolateral prefrontal cortex (rDLPFC) and right posterior parietal cortex (rPPC), and the underlying neurocognitive mechanisms, remain unclear. To determine the specific contributions of these regions, we applied continuous theta burst stimulation (cTBS) to the rDLPFC or rPPC in separate groups of non-clinical alcohol users (rDLPFC: n = 29; rPPC: n = 28) using a within-subject, active-sham counterbalanced design. Alcohol approach tendencies were assessed using the Alcohol Approach-Avoidance Task at baseline and following active and sham stimulation. Active rDLPFC cTBS selectively slowed alcohol push responses without affecting pull responses, whereas rPPC cTBS produced a bidirectional, alcohol-specific shift in which push responses slowed and pull responses simultaneously accelerated. Although stimulation at either site shifted automatic tendencies toward greater alcohol approach, the behavioral profiles were dissociable. The findings suggest that rDLPFC contributes to sustaining the regulatory signal that drives avoidance over prepotent alcohol approach responses, whereas rPPC determines whether that signal is successfully expressed in behavior when competing cue-driven influences are present. These findings provide node-specific evidence for functional specialization within the frontoparietal network and identify a potential parietal mechanism linking cue salience to overt alcohol approach behavior.

## Introduction

Automatic alcohol approach tendencies are one of the major determinants of harmful alcohol use and have been consistently associated with relapse risk and treatment outcomes (Barkby et al., 2012; Bowley et al., 2013; Field et al., 2017; Quoilin et al., 2021; Wiers et al., 2011). These tendencies reflect a state where alcohol-related cues acquire heightened incentive salience, as proposed by the incentive sensitization theory (Robinson & Berridge, 1993, 2025), and preferentially elicit approach responses before deliberate evaluation can intervene. Such cue-driven influences become increasingly dominant as drinking patterns consolidate, with progressive weakening of voluntary control over behavior (Fleming & Bartholow, 2014; Giannone et al., 2024).

The impaired Response Inhibition and Salience Attribution (iRISA) model suggests that automatic responses to alcohol cues arise from the combined influence of heightened cue salience and diminished executive control (Ceceli et al., 2026; Goldstein & Volkow, 2002). According to this framework, salience attribution increases the motivational priority of alcohol-related cues, biasing behavior toward approach, while inhibitory control suppresses these cue-driven responses when they conflict with current goals. The behavioral outcome is therefore determined by the interaction between the two processes. Although substantial evidence supports the involvement of salience attribution and inhibitory control in alcohol-related behavior (Ceceli et al., 2026; Goldstein & Volkow, 2002, 2011; Lindgren et al., 2019; McClure & Bickel, 2014), how these processes are implemented within the brain to regulate automatic alcohol approach tendencies remains unclear.

Neurocognitive evidence has consistently linked right frontal regions to the control of behavior under conditions of cognitive conflict (Cole et al., 2014; McNeill et al., 2018; Miller & Cohen, 2001). Among these regions, the right dorsolateral prefrontal cortex (rDLPFC) plays a central role in maintaining task goals and regulating behavior when prepotent responses conflict with current intentions (Ehlis et al., 2019, 2024). Disruption of the rDLPFC activity has been observed to be associated with reduced inhibitory control in the presence of alcohol cues (McNeill et al., 2018), whereas enhanced rDLPFC engagement has been associated with more effective suppression of cue-driven approach responses in favor of task requirements (Ernst et al., 2014; Verma et al., 2026), supporting its role in maintaining goal-directed control over automatic alcohol-related behavior.

In addition to frontal regions, parietal regions may also contribute to the regulation of prepotent responses (Verma et al., 2025; Wetherill et al., 2012). Accumulating evidence suggests that, beyond directing attention, the rPPC helps implement the selected action in the presence of motivationally salient cues (Grent-‘t-Jong & Woldorff, 2007; Hilgetag et al., 2001; Sengupta et al., 2024; Thut et al., 2005). In the context of alcohol cues, disruption of rPPC functioning may therefore impair the implementation of the selected action, allowing the salience-driven approach tendency to dominate behavioral output. Supporting this account, reduced rPPC activity has been reported to be associated with stronger alcohol approach tendencies (Verma et al., 2025).

While converging evidence implicates the rDLPFC and rPPC in alcohol approach tendencies, their respective causal contributions have not been directly examined. The present study applies continuous theta burst stimulation (cTBS) to the rDLPFC or rPPC in separate groups to examine the causal contributions of these regions to automatic alcohol approach tendencies, assessed using the Alcohol Approach-Avoidance Task (A-AAT; Kersbergen et al., 2015). cTBS has traditionally been associated with transient suppression of cortical excitability at the stimulated site (Huang et al., 2005; Tupak et al., 2013), but accumulating evidence indicates considerable variability in both the direction and magnitude of its effects across individuals (Brownjohn et al., 2014; McCalley et al., 2021). Accordingly, the study hypotheses were formulated based on the conventional inhibitory characterization of cTBS, while the functional consequences of stimulation were independently assessed using tasks sensitive to the cognitive functions associated with each region (see Methods).

Based on the proposed roles of rDLPFC and rPPC in cognitive processes relevant to alcohol approach tendencies, rDLPFC cTBS was expected to increase automatic approach tendencies by impairing inhibitory control over prepotent responses. rPPC cTBS was hypothesized to weaken the regulatory balance between required actions and cue salience, particularly during incongruent alcohol avoidance responses. Given reported individual differences in automatic alcohol tendencies, whereby some individuals exhibit automatic approach, and others automatic avoidance despite continued alcohol use (Barkby et al., 2012; Cousijn et al., 2014; Morris et al., 2020; Verma et al., 2025), baseline automatic tendency was also included as an exploratory factor.

## Methods

### Participants

An a priori power analysis using G*Power assuming a medium effect size (Cohen’s *f* = 0.25), α = 0.05, and power = 0.80 indicated a requirement of 28 participants per group (56 total) for a two-group, repeated-measurement design (Faul et al., 2007). Adjusting for an anticipated 10% attrition rate, the target sample was set to 62 participants (31 per group). Four participants were subsequently excluded: three from the rPPC group and one from the rDLPFC group withdrew. One additional participant in the rDLPFC group was excluded because the resting motor threshold could not be established due to failure to localize M1. The final sample comprised 29 participants in the rDLPFC group (mean age = 23.3 ± 2.94 years; 15 female) and 28 participants in the rPPC group (mean age = 23.0 ± 2.81 years; 13 female). All participants were active alcohol users with normal or corrected-to-normal vision, no history of neurological or psychiatric conditions, and were right- or mixed-right-handed. TMS eligibility was assessed using the Standard Screening Questionnaire for rTMS Candidates (Rossi et al., 2011). Alcohol use severity was assessed with the Alcohol Use Disorders Identification Test (AUDIT) (Saunders et al., 1993); no participant met diagnostic criteria for alcohol dependence at enrolment. The study was approved by the Institutional Ethics Committees of the University of Hyderabad (UH/IEC/2026/006) and the Indian Institute of Technology Hyderabad (IITH/IEC/2025/01/13) and conducted in accordance with the Declaration of Helsinki. All participants provided written informed consent prior to participation.

## Tasks and Measures

### Alcohol Approach–Avoidance Task (A-AAT)

Automatic approach and avoidance tendencies toward alcohol cues were assessed using an adapted A-AAT (Kersbergen et al., 2015; Verma et al., 2025) (Figure 1A and 1B). The task comprised 190 trials: two experimental blocks of 80 trials each, two practice blocks of 5 trials each, and a 20-trial washout block between the experimental blocks. Experimental trials contained equal proportions of alcohol-related and non-alcohol beverage images, pseudorandomized with a constraint against more than three consecutive same-category presentations. In the first experimental block, participants pulled the joystick for alcohol images and pushed for non-alcohol images; mappings were reversed in the second block. The washout block used neutral object images with the same push–pull responses to attenuate carry-over and use-dependent motor effects. Each trial began with a 500 ms fixation, followed by up to 1500 ms to respond. A minimum 200 ms neutral hold prior to stimulus onset was enforced to avoid premature responses. Pull movements produced a continuous zoom-in effect and push movements a zoom-out effect, scaled to joystick displacement along the y-axis. Feedback on incorrect responses was provided during experimental trials; full feedback (correct and incorrect) was given during practice. The task was administered across all three sessions, i.e., baseline, active, and sham.

**Figure 1.**
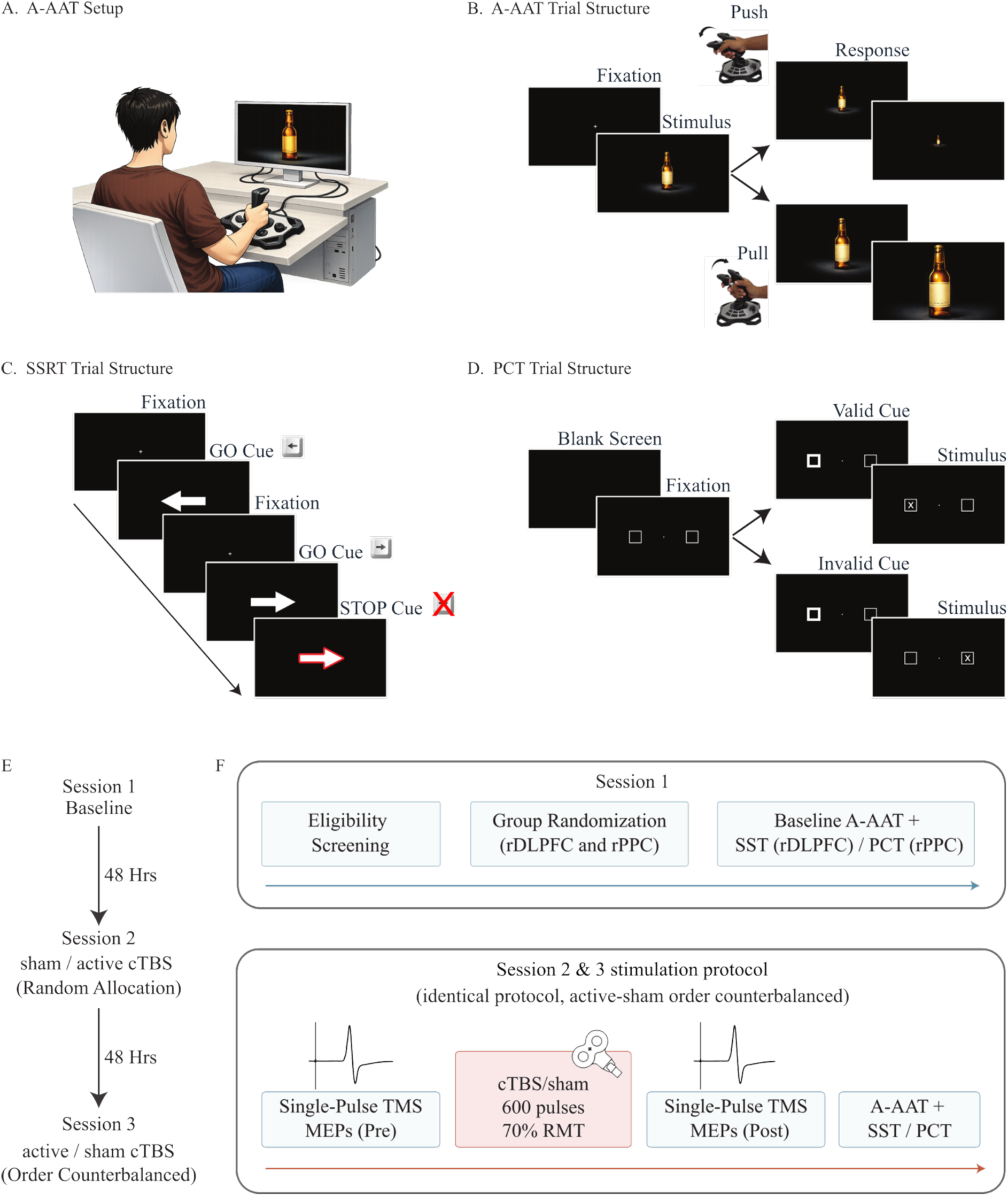
Experimental setup, A-AAT, SSRT, and PCT task structures, and experimental timeline for the cTBS investigation. Panel A illustrates the A-AAT setup, in which stimuli are presented on a landscape-oriented monitor and participants respond using a joystick with push and pull movements along the y-axis. Panel B depicts the A-AAT trial structure: each trial begins with a fixation, followed by stimulus onset, after which participants respond using the joystick. Pull responses produce a zoom-in effect, whereas push responses produce a zoom-out effect, both proportional to the joystick’s y-axis displacement, until the stimulus disappears at ±1. Panel C shows the Stop-Signal Task (SST) trial structure. Each trial begins with a fixation, followed by the presentation of a left or right arrow. On approximately 25% of trials, the arrow outline turns red, indicating a stop signal, and participants are required to inhibit their response. Panel D illustrates the Posner Cueing Task (PCT) trial structure. Each trial begins with a blank screen, followed by a fixation display with peripheral placeholders on the left and right. One placeholder briefly highlights, after which a target (“X”) appears either in the same location (valid trial) or the opposite location (invalid trial). Panel E illustrates the timeline of the three experimental sessions, each separated by a 48-hour interval to minimize practice effects and carryover effects of magnetic brain stimulation. In Session 1, participants were screened for active alcohol consumption and TMS compatibility, after which they were randomly assigned to either the rDLPFC or rPPC stimulation group (Panel F). Following group allocation, a baseline AAT was administered to classify participants based on automatic tendencies. Participants then completed 32 practice trials of the SST (rDLPFC group) or PCT (rPPC group). In Session 2, participants in each group were randomly assigned to receive either sham or active stimulation. Resting motor threshold (RMT) was determined, and the stimulation site was localized, followed by the recording of 20 pre-cTBS MEPs to assess baseline motor cortex excitability. cTBS consisted of triplets of pulses at 50 Hz delivered at 5 Hz continuously for 40 s (600 pulses total) at 70% RMT, followed by 20 post-cTBS MEP recordings. After an interval of approximately 10 minutes, participants completed the alcohol approach-avoidance task (A-AAT) (190 trials) and the SST or PCT (128 trials). Session 3 followed the same procedure as Session 2, with the exception that the stimulation condition (sham vs. active) was counterbalanced relative to Session 2. **Note.** The stimulus-display proportions shown in the figure are adjusted for visualization and do not reflect the exact dimensions used in the task.

### Functional Checks on Stimulation Efficacy

cTBS effects on cortical excitability can vary across individuals and sites, making it important to verify target engagement and functional modulation at the sites before interpreting its effects on alcohol approach-avoidance behavior. Participants in the rDLPFC and rPPC stimulation group completed the Stop-Signal Task and Posner Cueing Task, respectively. Stop-signal reaction time (SSRT) from SST provides a well-established measure of inhibitory control and the efficiency of the stopping process (Logan & Cowan, 1984; McNeill et al., 2018), whereas cueing cost on the PCT indexes attentional reorienting and the ability to disengage and reallocate attentional priority (Hayward & Ristic, 2013; Posner, 1980). Under the conventional inhibitory characterization of cTBS, stimulation of rDLPFC was expected to increase SSRT, reflecting reduced efficiency of the frontal inhibitory control process, whereas stimulation of rPPC was expected to increase attentional reorienting cost, reflecting impaired disengagement and reallocation of attentional priority. These measures were used to verify functional perturbation of selected regions.

#### Stop-Signal Task (SST)

The SST was administered to the rDLPFC group to assess whether cTBS modulated prefrontal inhibitory control (Logan & Cowan, 1984; McNeill et al., 2018). Participants responded to a centrally presented left- or right-pointing arrow by pressing the corresponding key (Figure 1C). Go trials constituted 75% of trials; on stop trials (25%), the arrow turned red after a variable stop-signal delay (SSD), cueing response inhibition. SSD was adjusted dynamically using a one-up/one-down staircase (Logan & Cowan, 1984), incrementing by 50 ms after successful inhibition and decrementing by 50 ms after failed inhibition, constrained between 50 and 500 ms (McNeill et al., 2018), targeting an inhibition success rate of approximately 50%. Each trial began with a 250 ms fixation cross, followed by the go stimulus with a 2000 ms response window, and a variable inter-trial interval of 500–1500 ms.

#### Posner Cueing Task (PCT)

The PCT was administered to the rPPC group to assess whether cTBS modulated parietal attentional processes. Attentional orienting was assessed using an exogenous spatial cueing paradigm (Hayward & Ristic, 2013; Posner, 1980). Each trial began with a blank screen for 250 ms, followed by a central fixation point flanked by two peripheral placeholder boxes for 750 ms (Figure 1D). A peripheral cue was presented for 80 ms to induce automatic attentional capture. Cue validity was manipulated such that cues were valid on 60% of trials, invalid on 20%, and neutral on 20%. Cue–target intervals were varied across early, intermediate, and late temporal windows to sample multiple phases of attentional orienting. Following the cue–target interval, the letter “X” appeared in one placeholder box; participants responded to the target onset by pressing the spacebar as quickly as possible, within a 2000-ms window. Trials concluded with a 750 ms fixed inter-trial interval.

### Theta Burst Stimulation Procedure

TMS was delivered using a DuoMAG XT100 stimulator (Deymed Diagnostics, Czech Republic) with a BF70 figure-of-eight coil. Stimulation sites were localized prior to threshold determination. The left M1 hand area was identified using a frameless stereotaxic neuronavigation system (Brainsight v2.0; NDI Polaris Vicra) by targeting the hand knob region of the precentral gyrus on the MNI template, with coil position iteratively adjusted in ∼1 cm increments until stimulation reliably evoked motor-evoked potentials (MEPs) or visible contractions of the right first dorsal interosseous (FDI) muscle. In the rDLPFC group, the F4 electrode location was identified using the international 10–20 EEG system combined with the Beam-F4 method as the stimulation target (Beam et al., 2009). In the rPPC group, the P4 location was identified using the 10–20 system. Resting motor threshold (RMT) was determined at the start of each session as the minimum stimulator intensity evoking peak-to-peak MEPs of ≥50 μV in the relaxed right FDI in at least 5 of 10 consecutive trials. EMG was recorded via Ag–AgCl surface electrodes in a belly–tendon montage, with the reference electrode over the ulnar bone. The cTBS protocol comprised bursts of three pulses at 50 Hz repeated at 5 Hz, applied continuously for 40 s (600 pulses total; Huang et al., 2005), at 70% RMT, administered 10 minutes before task performance. Sham stimulation was delivered using the 45° tilted-coil procedure (Wang et al., 2024), preserving auditory and somatosensory characteristics of active stimulation while minimizing cortical activation.

### Experimental Procedure

A within-subject repeated-measures design was employed, with all participants completing three sessions separated by 48-hour intervals to minimize practice and carry-over effects of cTBS (Schwippel et al., 2025) (Figure 1E). Session 1 involved eligibility screening, informed consent, and baseline assessment using the alcohol approach–avoidance task (A-AAT) to classify participants based on their approach or avoidance tendencies toward alcohol cues (Figure 1F). Participants were then randomly assigned to either the rDLPFC or rPPC stimulation group. Following group allocation, participants completed 32 practice trials of the Stop-Signal Task (SST; rDLPFC group) or the Posner Cueing Task (PCT; rPPC group) for task familiarization. The SST was administered as a functional check on rDLPFC modulation, given that SSRT is sensitive to frontal disruption (Chen et al., 2021; McNeill et al., 2018), and an increase following active cTBS would be consistent with modulation of the inhibitory control processes associated with the targeted region. The PCT was administered as a functional check on rPPC modulation, as attentional reorienting cost provides a sensitive index of parietal function and is known to be affected by parietal disruption (Capotosto et al., 2012; Sengupta et al., 2024).

Sessions 2 and 3 followed an identical structure, with the exception that the stimulation condition (active vs. sham cTBS) was counterbalanced across sessions (Figure 1F). In each session, resting motor threshold (RMT) was first determined, and the stimulation site was localized. Motor evoked potentials (MEPs) were then recorded from the primary motor cortex (M1) prior to stimulation (20 pulses) to assess baseline corticospinal excitability. Continuous theta burst stimulation (cTBS) was delivered as triplets of pulses at 50 Hz repeated at 5 Hz for 40 s (600 pulses total) at 70% RMT. Immediately following stimulation, 20 post-cTBS MEPs were recorded to assess changes in motor cortex excitability. After an interval of approximately 10 minutes, participants completed the A-AAT (190 trials) as the primary outcome measure. Subsequently, participants performed either the SST (rDLPFC group) or the PCT (rPPC group), consisting of 128 trials administered in two blocks of 64 trials separated by a 30 s rest period, to assess cTBS’ effect on inhibitory control and attentional orienting, respectively.

### Data Analysis

#### Alcohol Approach–Avoidance Index

Response time (RT) in the A-AAT was defined as the interval from stimulus onset to maximum joystick displacement (y = ±1). The alcohol approach index (AAI) was calculated as: (RT_push alcohol_ − RT_pull alcohol_) − (RT_push non-alcohol_ − RT_pull non-alcohol_), with higher values indicating stronger approach tendencies toward alcohol cues. Because classification of participants by automatic tendency (approach vs. avoidance) was derived post hoc, analyses involving this factor were treated as exploratory.

#### Stop-Signal Reaction Time

Stop-signal reaction time (SSRT) was estimated using the integration method (McNeill et al., 2018; Verbruggen et al., 2019). Anticipatory responses (RT < 100 ms) and trials with no valid response were excluded. Go trial RTs were rank-ordered, and the n^th^ RT corresponding to the proportion of failed inhibitions was identified, from which the mean SSD was subtracted to yield SSRT. SSRT estimates were accepted only when the probability of successful inhibition fell between 0.25 and 0.75, and the mean RT on failed stop trials was shorter than the mean go RT, per the horse-race model assumptions (Logan & Cowan, 1984; Verbruggen et al., 2019).

#### Attentional Orienting Cost

Mean RTs were computed separately for valid and invalid cue trials. Reorienting cost was defined as RT_invalid_ − RT_valid_, indexing the cost of disengaging from an invalid cue. Higher reorienting cost indicates less efficient attentional reorienting.

#### Statistical Analysis

The effects of cTBS on inhibitory control (SSRT; rDLPFC group), attentional orienting (reorienting cost; rPPC group), and AAI were examined using repeated-measures ANOVA with Stimulation (active vs. sham) and Automatic Tendency (approach vs. avoidance) as factors, with stimulation order included as a covariate. Significant effects were followed up with paired-samples t-tests to characterize the direction of differences. To identify the components contributing to changes in AAI, reaction times for each condition (alcohol pull, alcohol push, non-alcohol pull, and non-alcohol push) were analyzed separately, with comparisons made between active and sham stimulation conditions. To assess potential stimulation order (sham vs active) effects, differences in AAI change scores were compared between order conditions using independent-samples *t*-tests within each group, with Welch’s correction applied where appropriate (i.e., unequal variances), resulting in fractional degrees of freedom. Three participants did not yield valid SSRT estimates in the sham condition because the probability of successful inhibition on stop trials exceeded the recommended range (i.e., > 0.75; Verbruggen et al., 2019). Consequently, paired SSRT analyses were conducted on complete cases only (*n* = 26). The change in motor cortex excitability post-cTBS was assessed using paired Wilcoxon signed-rank tests for pre–post comparisons, due to violations of normality, and supplemented by analyses of normalized (% baseline) data tested against 100 using a one-sample Wilcoxon test. P-values were adjusted for multiple comparisons using the Benjamini-Hochberg procedure. All statistical analyses were conducted using R (version 4.4.1; R Core Team, 2024).

## Results

### Participant Characteristics

During session 1, the baseline A-AAT was administered to assess participants’ pre-existing automatic tendencies toward alcohol cues. Participants were classified post-hoc based on baseline AAI scores, with positive values indicating approach and negative values indicating avoidance tendencies. This classification was included as an exploratory factor in subsequent analyses to examine whether cTBS effects on alcohol approach–avoidance behavior were moderated by individuals’ baseline automatic tendencies. The distribution of participants across automatic tendencies was unequal (Table 1), with fewer avoidance tendency individuals across groups, which reduced statistical power to detect tendency-based differences. Within the rDLPFC group, approach and avoidance individuals did not differ significantly in age (*t_(7.04)_* = −0.87, *p* = 0.413), AUDIT scores (*t_(9.42)_* = 1.07, *p* = 0.312), AUDIT risk category (*p* = 1.00), or education level (*p* = 0.625). Similarly, within the rPPC group, approach and avoidance individuals did not differ in age (*t_(23.71)_* = 1.34, *p* = 0.194), AUDIT scores (*t_(25.43)_* = 0.66, *p* = 0.517), risk category (*p* = 1.00), and education (*p* = 0.682). Across the combined sample, baseline AAI was not significantly correlated with AUDIT scores (*r* = 0.24, *p* = 0.078), suggesting that baseline automatic alcohol tendencies varied independently of alcohol-use severity. Approach and avoidance individuals were thus comparable on demographic and alcohol-use characteristics within both stimulation groups.

**Table 1.** Demographics and baseline characteristics of participants across stimulation groups and automatic tendencies.

|  | rDLPFC |  | rPPC |  |
| --- | --- | --- | --- | --- |
|  | Approach<br>(N=23) | Avoidance<br>(N=6) | Approach<br>(N=18) | Avoidance<br>(N=10) |
| Age (mean ± SD) | 23.0 ± 2.85 | 24.3 ± 3.33 | 23.4 ± 3.03 | 22.1 ± 2.23 |
| AAI (Mean ± SE) | 0.091 ± 0.015 | -0.042 ± 0.012 | 0.070 ± 0.014 | -0.043 ± 0.012 |
| AUDIT Score (Mean ± SE) | 5.13 ± 0.56 | 4 ± 0.89 | 6 ± 0.907 | 5.3 ± 0.559 |
| AUDIT risk category |  |  |  |  |
| Low risk (1-7) (n) | 20 (86.96%) | 5 (83.33%) | 14 (77.18%) | 9 (90.00%) |
| Moderate risk (8-14) (n) | 3 (13.04%) | 0 (0.00%) | 3 (16.67%) | 1 (10.00%) |
| High risk (15+) (n) | 0 (0.00%) | 1 (16.67%) | 0 (0.00%) | 0 (0.00%) |
| Education |  |  |  |  |
| Undergraduate (n) | 8 (34.78%) | 1 (16.67%) | 5 (27.78%) | 4 (40.00%) |
| Postgraduate (n) | 9 (39.13%) | 2 (33.33%) | 8 (44.44%) | 5 (50.00%) |
| Doctorate (n) | 6 (26.09%) | 3 (50.00%) | 5 (27.78%) | 1 (10.00%) |

*Note.* AAI = Alcohol Approach Index; AUDIT = Alcohol Use Disorder Identification Test.

### Verification of cTBS-Induced Cognitive Modulation

Initially, potential order effects (active vs. sham cTBS) across the second and third sessions were assessed for the primary outcome variable (AAI), where no significant stimulation order effects were observed. In the rPPC group, the change in AAI (ΔAAI) did not differ between stimulation order conditions (active-before-sham vs. active-after-sham), *t_(22.60)_* = 0.08, *p* = 0.935, *95% CI* [−0.062, 0.067]. Mean ΔAAI was comparable for participants receiving active stimulation before sham (*M* = 0.053) and those receiving it after sham (*M* = 0.051). Similarly, in the rDLPFC group, ΔAAI did not significantly differ as a function of stimulation order, *t_(26.61)_* = 0.68, *p* = 0.500, *95% CI* [−0.048, 0.095], with a mean ΔAAI of 0.057 for the active-before-sham condition and 0.033 for the active-after-sham condition.

To verify that active cTBS modulated the targeted cognitive processes at each stimulation site, inhibitory control was assessed via SSRT in the rDLPFC group and attentional reorienting via PCT reorienting cost in the rPPC group. In the rDLPFC group, a main effect of stimulation was observed for SSRT (*F_(1,23_)* = 10.13, *p* = 0.004, *η²_p_* = 0.306), with a significant increase in SSRT after active stimulation relative to sham (*t_(25)_* = −4.46, *p* < 0.001, *d* = 0.88, *ΔM* = 0.056; Figure 2A). This pattern was consistent with disruption of the prefrontal inhibitory control mechanism indexed by SSRT. Neither the stimulation × order interaction (*F_(1,23)_* = 0.046, p = 0.832) nor the stimulation × tendency interaction (*F_(1,23)_* = 0.049, p = 0.827) was significant, indicating that the effect of stimulation was consistent across stimulation order and baseline automatic tendencies.

**Figure 2.**
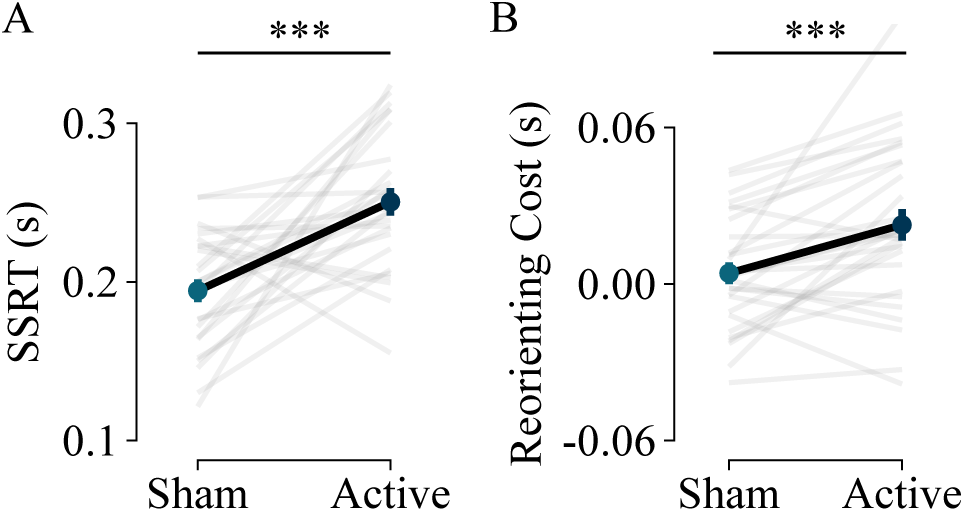
In the rDLPFC group, SSRT was significantly increased following active cTBS compared to sham stimulation, indicating reduced inhibitory control (Panel A). Similarly, in the rPPC group, reorienting cost in the PCT was significantly increased after active cTBS, reflecting impaired attentional reorienting (Panel B). Grey lines depict paired data for individual participants, while black lines represent group means under sham and active stimulation conditions. *** p < 0.001.

Similarly, in the rPPC group, a main effect of stimulation was observed for PCT reorienting cost (*F_(1,25)_* = 14.17, *p* < 0.001, *η²_p_* = 0.362), where active stimulation significantly increased attentional reorienting cost relative to sham (*t_(27)_* = −3.85, *p* = 0.001, *d* = 0.73, *ΔM* = 0.019), consistent with disruption of the parietal priority-disengagement mechanism (Figure 2B). Further, no significant stimulation × order (*F_(1,25)_* = 2.70, *p* = 0.113) or the stimulation × tendency interaction (*F_(1,25)_* = 0.134, *p* = 0.717) was observed in rPPC group. These results align with the conventional inhibitory characterization of cTBS and suggest successful modulation of the targeted cognitive processes in both groups.

Additionally, because the A-AAT involves different motor responses depending on stimulus type and experimental block, motor cortex excitability was assessed as a control measure. Representative MEP waveforms (Figure 3A and 3C) illustrate comparable pre- and post-stimulation responses across conditions and groups. In the rDLPFC group, MEP amplitudes did not significantly differ between pre- and post-stimulation phases following active stimulation (*ΔMdn* = 25.5, *V* = 114, *p* = 0.123) or sham stimulation (*ΔMdn* = −8.54, *V* = 186, *p* = 0.803). Similarly, in the rPPC group, no significant pre–post changes were observed following active (*ΔMdn* = 22.5, *V* = 121, *p* = 0.173) or sham stimulation (*ΔMdn* = −3.52, *V* = 212, *p* = 0.367). To account for baseline variability, the percentage change from baseline also showed no significant deviation from baseline (100%) in any condition (all *p*s > 0.05; Figure 3B and 3D). These results indicate that general motor cortex excitability was not altered by stimulation at either site, suggesting that the behavioral effects observed during the A-AAT were not attributable to non-specific changes in motor output.

**Figure 3.**
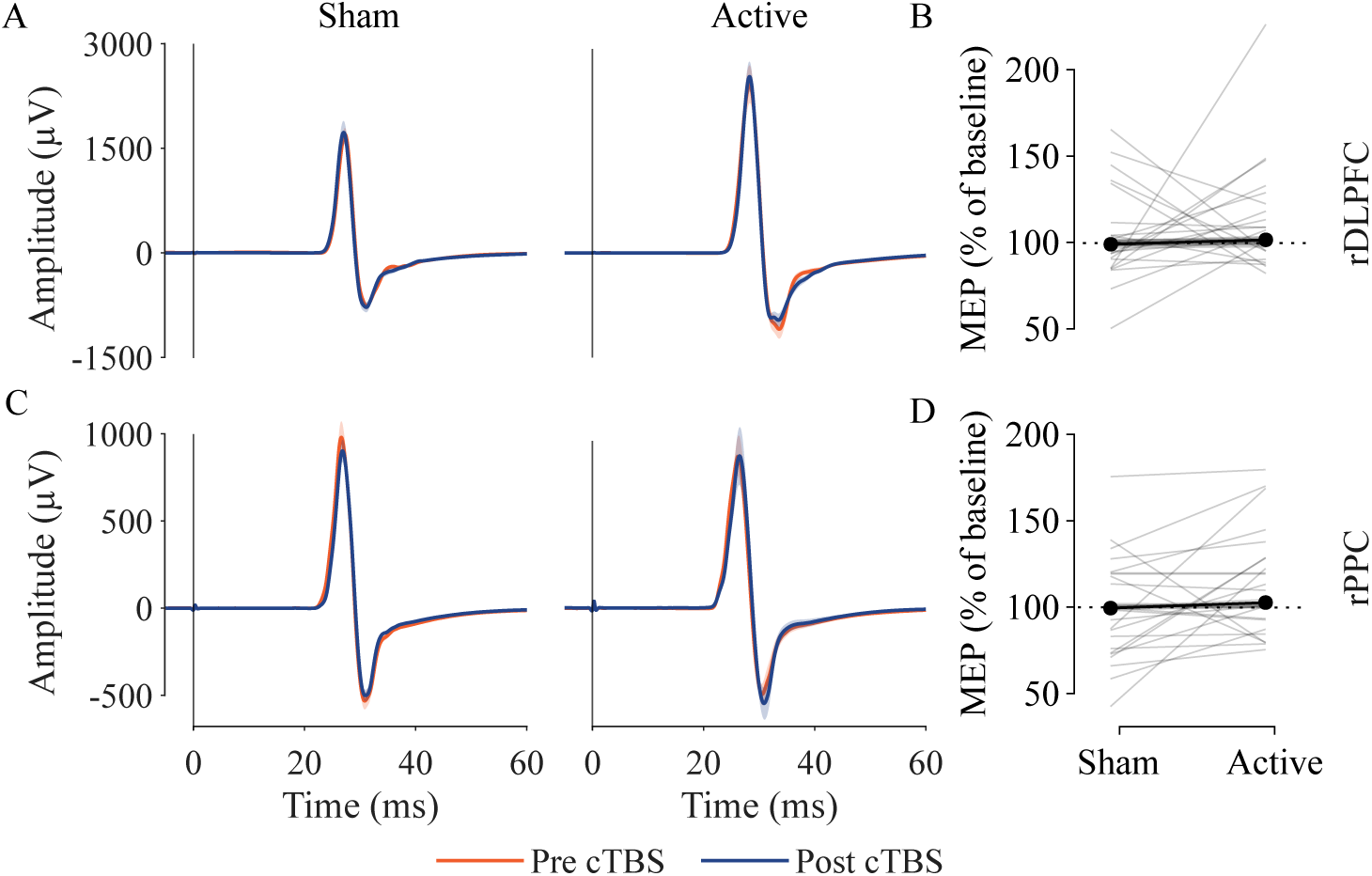
Panels A and C show representative MEP waveforms from a single participant (FDI muscle; single-pulse TMS at 120% RMT) in the rDLPFC and rPPC conditions, before and after sham and active stimulation. Each waveform reflects the average of 20 MEPs and is shown for illustration, with red traces representing pre-cTBS MEP waveforms and blue traces representing post-cTBS MEP waveforms. Panels B and D present group-level results, where percent change relative to pre-cTBS baseline revealed no significant change in either the rDLPFC (Panel B) or rPPC (Panel D) groups. Grey lines represent paired data from individual participants, while black lines indicate group means under sham and active stimulation conditions.

### Effects of cTBS on Alcohol Approach–Avoidance Tendencies

To examine the behavioral consequences of cTBS at rDLPFC and rPPC locations on alcohol approach–avoidance behavior, AAI was compared following active and sham cTBS, while treating stimulation order as a covariate. Active stimulation produced a significant influence on AAI in both groups (rDLPFC: *F_(1, 26)_* = 4.41, *p* = 0.045, *η²_p_* = 0.145; rPPC: *F_(1, 25)_* = 5.51, *p* = 0.027, *η²_p_* = 0.181), where stimulation at either region shifted automatic tendencies toward greater alcohol approach (rDLPFC: *t_(28)_* = −2.52, *p* = 0.018, *d* = 0.47, *ΔM* = 0.045, Figure 4A; rPPC: *t_(27)_* = −3.41, *p* = 0.002, *d* = 0.64, *ΔM* = 0.052, Figure 4C). The stimulation × tendency interaction was not significant in either group (rDLPFC: *F_(1, 26)_* = 0.006, *p* = 0.940; rPPC: *F_(1, 25)_* = 0.011, *p* = 0.918). No main effect of stimulation (active vs sham) order was found across regions, or its interaction with other factors (all ps > 0.05).

**Figure 4.**
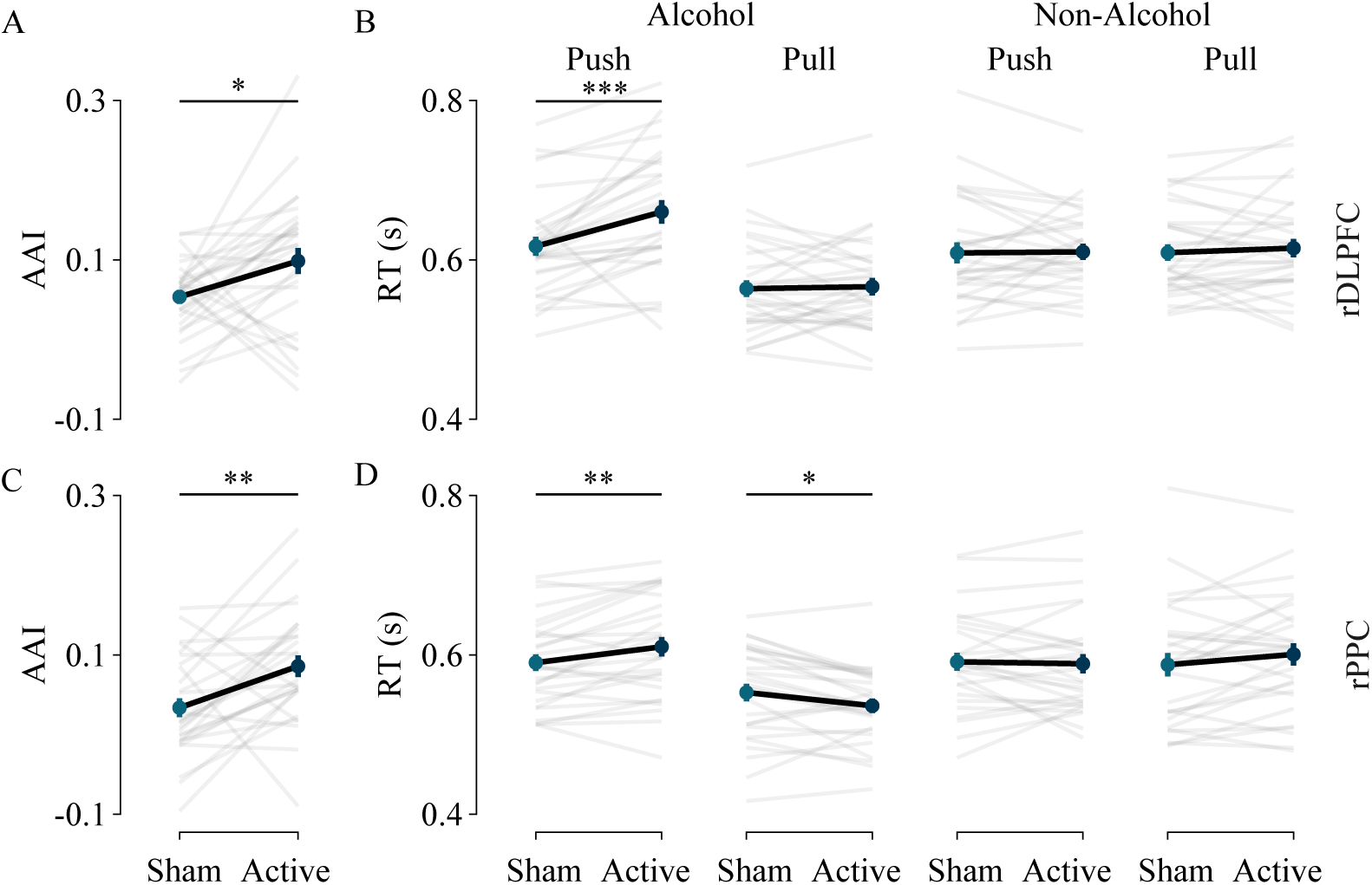
Active cTBS to the rDLPFC significantly increased AAI (Panel A). This effect was selectively driven by slower responses during alcohol push trials (Panel B), with no corresponding changes in pull or non-alcohol conditions, suggesting a disruption of avoidance-specific processes. Active cTBS to the rPPC also resulted in an overall increase in AAI (Panel C), primarily driven by a bidirectional, alcohol-specific effect characterized by slower alcohol push responses and faster alcohol pull responses (Panel D), with no changes observed in non-alcohol conditions. Grey lines represent paired data from individual participants, while black lines indicate group means under sham and active stimulation conditions. * p < 0.05; ** p < 0.01; *** p < 0.001.

Analyses excluding participants with baseline avoidance tendencies produced the same overall pattern of stimulation effects in both groups (rDLPFC: *F_(1,21)_* = 4.51, *p* = 0.046, *η²_p_* = 0.177; rPPC: *F_(1,16)_* = 6.10, *p* = 0.025, *η²_p_* = 0.276), indicating that the primary findings were not dependent on inclusion of the smaller avoidance subgroup. Accordingly, all participants, including those with baseline avoidance tendencies, were included in the subsequent analyses to maximize statistical power. However, this should not be interpreted as evidence that baseline avoidance tendency has no influence on stimulation effects. Rather, given the substantial imbalance in subgroup sizes, the study was underpowered to reliably detect moderation by baseline tendency, and the absence of a significant interaction should therefore be interpreted with caution.

### Source of Increased Alcohol Approach Tendencies

To identify the behavioral source of changes in alcohol approach tendencies, individual AAI components were examined separately. In the rDLPFC group, active stimulation selectively slowed alcohol push responses relative to sham (*t_(28)_* = −4.20, *p* < 0.001, *d* = 0.78, *ΔM* = 0.043) with no significant effects on alcohol pull (*t_(28)_* = −0.31, *p* < 0.759, *ΔM* = 0.003; Figure 4B). In the rPPC group, active stimulation significantly slowed alcohol push responses (*t_(27)_* = −3.35, *p* = 0.002, *d* = 0.63, *ΔM* = 0.020), while alcohol pull responses became significantly faster (*t_(27)_* = 2.74, *p* = 0.011, *d* = 0.52, *ΔM* = −0.017, Figure 4D). Non-alcohol trials were unaffected by stimulation in both groups (all *p*s > 0.05, Figure 4B and 4D).

In sum, active cTBS modulated the targeted cognitive processes (response inhibition and attentional allocation), as evidenced by longer SSRT in the rDLPFC group and greater reorienting cost in the rPPC group, respectively. During the A-AAT, stimulation at both sites shifted behavior toward a greater alcohol approach through behaviorally dissociable pathways. cTBS at the rDLPFC site selectively slowed alcohol push responses without affecting pull, consistent with disruption of effortful avoidance control. cTBS at the rPPC site produced a bidirectional, alcohol-specific pattern (simultaneously slowing push and accelerating pull) consistent with disruption of competitive priority regulation over action representations. These dissociable profiles demonstrate a behavioral dissociation between the consequences of stimulating the two scalp locations within the same approach-avoidance paradigm.

## Discussion

The present study employed cTBS to examine the contributions of rDLPFC and rPPC to the behavioral expression of automatic alcohol approach tendencies. Although suppression of either site shifted behavior toward greater alcohol approach, the behavioral profiles associated with each stimulation site were dissociable. rDLPFC cTBS selectively slowed alcohol push responses, whereas rPPC cTBS both slowed alcohol push and accelerated alcohol pull responses. These dissociable profiles suggest that the two regions contribute to alcohol approach-avoidance regulation through partially distinct mechanisms, with rDLPFC generating and maintaining the control signal that drives avoidance, and rPPC translating the control signal into behavioral output in the presence of competing cue-driven influences.

### Prefrontal Regulation of Avoidance Signaling

The unidirectional effect of rDLPFC cTBS, with selectively slowed push responses and no change in pull responses in the presence of alcohol cues, suggests rDLPFC’s specific role in generating the signal that drives avoidance responses rather than suppressing approach responses (Cole et al., 2014; McNeill et al., 2018; Miller & Cohen, 2001). When this signal is weakened, avoidance slows, and the approach tendency may default in the absence of sufficient regulatory control. McNeill et al. (2018) demonstrated that cTBS to rDLPFC impairs inhibitory control and increases alcohol consumption (McNeill et al., 2018); the present findings extend this by suggesting such disruption may be caused by selective impairment of avoidance behavior in the presence of alcohol cues, rather than influencing approach responses.

Prior studies applying non-invasive stimulation to the left DLPFC have generally failed to produce reliable changes in alcohol approach behavior despite reporting changes in craving (Chase et al., 2011; den Uyl et al., 2015; Hanlon et al., 2015; McNeill et al., 2022; Paulus, 2007). This divergence between left and right hemispheric frontal regions may reflect functional specialization, with left frontal regions more commonly implicated in reward processing that later manifest into craving-related responses (den Uyl et al., 2015; McNeill et al., 2022), whereas right frontal regions may be more prominent in generating regulatory control over responses that support avoidance under conflict (Cole et al., 2014; McNeill et al., 2018; Miller & Cohen, 2001; Verma et al., 2025). Accordingly, avoidance responding may be specifically vulnerable to conditions that transiently reduce right prefrontal availability, such as acute stress, cognitive load, or intoxication, which are well-documented to increase relapse risk (Barkby et al., 2012; Wiers et al., 2011).

### Parietal Mediation of Avoidance Expression

The parietal cortex has primarily been implicated in attentional processes toward motivationally salient cues (Bisley & Goldberg, 2010; Grent-‘t-Jong & Woldorff, 2007; Sengupta et al., 2024; Thut et al., 2005), but the present findings suggest a more specific contribution extending beyond attentional allocation to the implementation of action when task demands conflict with competing cue-driven influences. Evidence from Contò et al. (2023) and Sengupta et al. (2024) suggests that rPPC regulates which action representations gain priority for behavioral output depending on task demands and cue salience (Contò et al., 2023; Sengupta et al., 2024), positioning it as a site where prefrontal control signals may be translated into overt behavior. When rPPC functioning was disrupted, avoidance responses slowed while approach responses simultaneously accelerated, suggesting that the translation of the avoidance control signal into behavior was impaired, allowing the competing cue salience-driven approach tendency to dominate behavioral output in its place.

### Integrated Understanding of Frontoparietal Contributions

Although the rDLPFC and rPPC findings come from separate groups, the evidence collectively suggests that rDLPFC generates the control signal that instructs avoidance, while rPPC mediates the behavioral expression of that signal in the presence of motivationally salient cues. When rPPC is impaired, the control signal may be present but fail to translate into the required action, allowing a cue-driven approach to prevail. Intact engagement of both regions may therefore be necessary to prevent dominant automatic approach tendencies from overriding controlled behavior, consistent with observations that reduced frontoparietal activity in alcohol users is associated with diminished capacity to regulate prepotent responses (Cofresí et al., 2026; van Oort et al., 2023).

These findings point to a mechanism not explicitly addressed within the iRISA framework (Ceceli et al., 2026; Goldstein & Volkow, 2002, 2011). Although the model emphasizes heightened salience attribution and diminished executive control as central features of addictive behavior, it provides limited consideration of the mechanisms through which salience signals are translated into overt action. The present results suggest that, beyond the frontal, orbitofrontal, and striatal systems emphasized by iRISA, the rPPC may contribute to this translational step by determining whether the control signal successfully drives avoidance behavior or is overridden by competing cue-driven approach tendencies at the point of action selection. The present findings may also provide a mechanistic explanation for the mixed efficacy of alcohol avoidance training reported in clinical trials (Manning et al., 2022; Spruyt et al., 2025). Given evidence that avoidance learning primarily recruits prefrontal mechanisms supporting goal-directed control (Ehret et al., 2024; Moscarello & LeDoux, 2013), interventions limited to enhancing these processes may be less effective if the parietal mechanisms responsible for implementing that control remain impaired.

From a translational perspective, these findings identify neural correlates that may be targeted through non-invasive brain stimulation, depending on where the impairment lies (frontal, parietal, or both). Whether enhancing activity within these regions would produce the inverse behavioral pattern remains to be established before clinical application. Future neuromodulation approaches may benefit from targeting these distinct mechanisms, with prefrontal approaches aimed at strengthening the regulatory control signal and parietal approaches aimed at improving the translation of that signal into required behavior in the presence of motivationally salient cues.

### Null Effect of Baseline Tendency

There was no significant interaction observed between stimulation and baseline tendency in either group, despite prior work demonstrating differential intervention outcomes depending on the magnitude of baseline automatic tendencies (Barkby et al., 2012; Morris et al., 2020; Verma et al., 2025). This null interaction is uninformative given the substantially unequal subgroup sizes resulting from post-hoc tendency-based classification, which limited power to detect moderation effects. Whether baseline tendency moderates the effects of frontoparietal stimulation, therefore, remains an open question for adequately powered future work.

## Limitations

Several limitations warrant acknowledgment in the present study. First, the sample comprised non-clinical active alcohol users rather than individuals with alcohol dependence. This was a deliberate choice to maximize mechanistic clarity, as clinical populations introduce confounds associated with chronic neuroadaptation, comorbidity, and dependence-related cognitive deficits (Fairbairn & Kang, 2025; Oscar-Berman & Marinković, 2007; Rehm et al., 2017; Topiwala et al., 2018). Whether disruption of rDLPFC and rPPC produces comparable behavioral consequences in individuals with established dependence cannot be determined from the present data. A second limitation concerns stimulation targeting and verification. Stimulation sites were localized using scalp-based procedures rather than individual MRI-guided neuronavigation, introducing uncertainty regarding the precise cortical regions affected. Although the SST and PCT provided functional evidence consistent with modulation of the targeted cognitive processes, no neurophysiological or neuroimaging measures were obtained to directly verify changes in neural activity. Furthermore, cTBS effects are not restricted to the stimulated site and may propagate through functionally connected networks, meaning the observed behavioral effects cannot be attributed exclusively to local modulation of rDLPFC or rPPC. The mechanistic account proposed here should therefore be viewed as a plausible interpretation consistent with the data rather than a directly verified causal explanation.

## Conclusion

The present study provides evidence for dissociable contributions of rDLPFC and rPPC to the behavioral expression of automatic alcohol approach tendencies, consistent with their roles at distinct stages of approach-avoidance regulation. Disruption of rDLPFC selectively impaired avoidance responding in the presence of alcohol cues without affecting approach, consistent with a role in generating the regulatory control signal that drives avoidance under motivational conflict. Disruption of rPPC produced a bidirectional, alcohol-specific shift, simultaneously impairing avoidance and facilitating approach, consistent with a role in translating that control signal into behavioral output when competing cue-driven influences are present. In particular, rPPC emerges as a node at which the avoidance signal may be overridden by motivationally salient cues, a contribution the iRISA model does not currently incorporate, while the rDLPFC findings extend the model’s account of executive control toward node and action-specific behavioral evidence. Together, these findings advance understanding of the frontoparietal architecture underlying automatic alcohol approach tendencies. The proposed mechanisms remain inferential and warrant further investigation through studies combining brain stimulation with neuroimaging and replication in clinical populations to establish their generalizability and clinical relevance.

## DECLARATIONS

## Data Availability

The datasets used and/or analyzed during the current study are available from the corresponding author upon reasonable request.

## Acknowledgment

This work was supported by the Ministry of Education, Government of India, through the Prime Minister’s Research Fellowship (PMRF) awarded to AKV.

